# Eco-evolutionary dynamics of pathogen immune-escape: deriving a population-level phylodynamic curve

**DOI:** 10.1101/2024.07.23.604819

**Authors:** Bjarke Frost Nielsen, Chadi M. Saad-Roy, C. Jessica E. Metcalf, Cécile Viboud, Bryan T. Grenfell

**Affiliations:** High Meadows Environmental Institute, Princeton University, Princeton, New Jersey, USA; Miller Institute for Basic Research in Science, University of California, Berkeley, California, USA; Department of Integrative Biology, University of California, Berkeley, California, USA; Department of Ecology and Evolutionary Biology, Princeton University, Princeton, New Jersey, USA; Division of International Epidemiology and Population Studies, Fogarty International Center, National Institutes of Health, Bethesda, Maryland, USA

## Abstract

The phylodynamic curve [1] conceptualizes how immunity shapes the rate of viral adaptation in a non-monotonic fashion, through its opposing effects on viral abundance and the strength of selection. However, concrete and quantitative model realizations of this influential concept are rare. Here, we present an analytic, stochastic framework in which a population-scale phylodynamic curve emerges dynamically, allowing us to address questions regarding the risk and timing of emergence of viral immune escape variants. We explore how pathogen- and population-specific parameters such as strength of immunity, transmissibility and antigenic constraints affect the phylodynamic curve, leading to distinct phylodynamic curves for different pathogens. Motivated by the COVID-19 pandemic, we probe the likely effects of non-pharmaceutical interventions (NPIs), and the lifting thereof, on the risk of viral escape variant emergence. Looking ahead, the framework has the potential to become a useful tool for probing how natural immunity, as well as choices in vaccine design and distribution and the implementation of NPIs affect the evolution of common viral pathogens.

## I. INTRODUCTION

The emergence of novel viral variants is widely understood to be a highly stochastic phenomenon [2, 3] and it follows that any statements concerning the risk of viral adaptation must be statistical in nature. This does not, however, detract from their usefulness or fundamental importance: the ability to pinpoint risk factors for the emergence of e.g. an immune escape variant is of great value to public health, especially if quantitative.

One example of such a statement regarding the risk of immune evasion at the within-host level is the phylodynamic curve (see Fig. 1A), introduced by [1]. While the concept has proved influential [3–7], there have been few quantitative realizations to date. [8] is a notable within-host exception and, importantly, [9, 10] address strain-replacement in influenza, focusing on the impact of strain-transcending immunity and complex fitness landscapes, respectively. In [11], the authors explored a model of the dynamics of antigenic distance (and thus immune escape) within a single influenza season. We argue that there is a further need for generic modeling frameworks to explore the relationship between individual and population-level epi-evolutionary dynamics. Aiming for a broadly applicable, reductionist framework addressing this gap, we develop an analytic description of the *net viral adaptation rate* (the immune escape variant emergence risk per unit of time) as shaped by population immunity (see Fig. 1B).

**FIG. 1.**
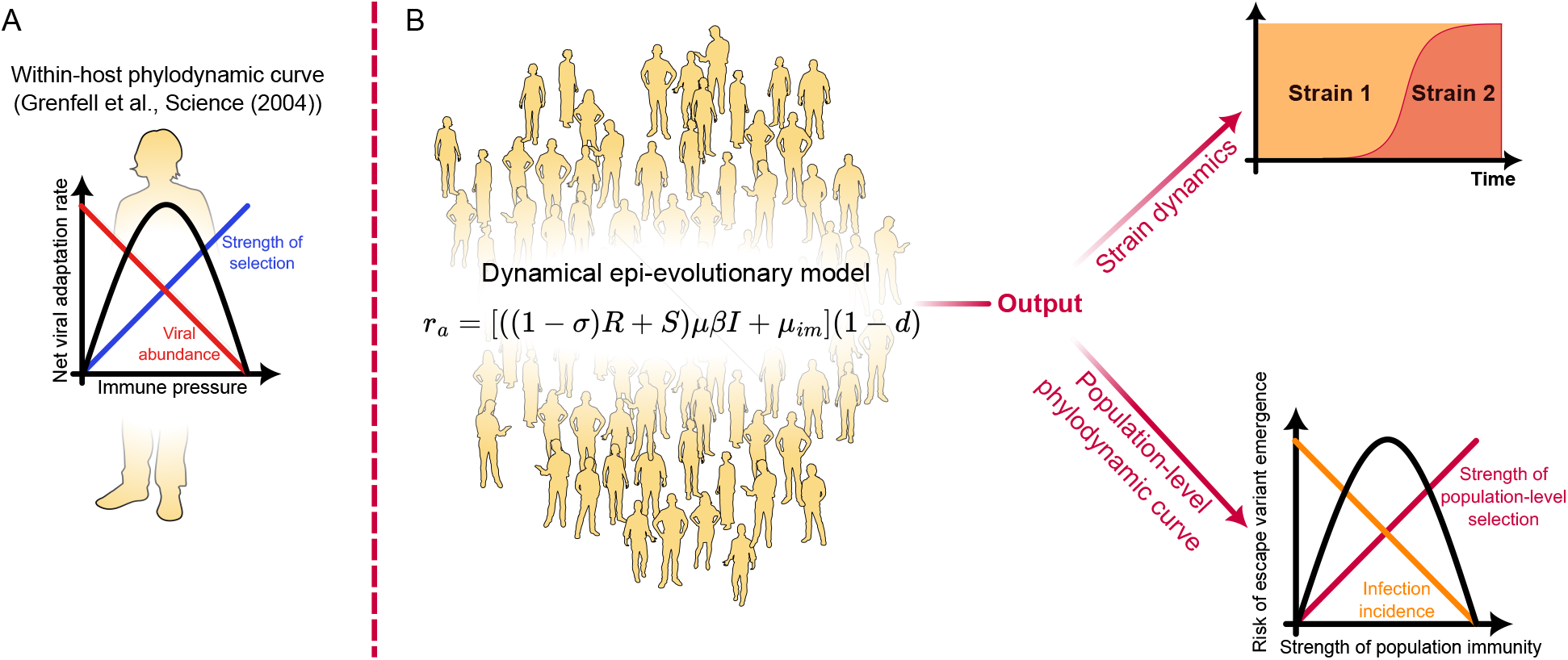
The model framework. **A)** The within-host phylodynamic curve, a concept introduced by [1]. When the immune response is sufficiently strong, abundance of virions is low and this limits viral adaptation. If the immune response is very weak or absent, there is little selection for immuneevasion, and thus adaptation is again limited. Peak adaptation is observed at intermediate strength of the immune response. **B)** A dynamical population-scale model is introduced, relying only on a few key assumptions about imperfect immunity, transmission and the occasional appearance of immune-evading mutations in infected hosts. From this framework, a population-scale analog of the phlodynamic curve dynamically emerges, addressing the risk of an immune escape variant arising and establishing in the population. Furthermore, the model allows exploration of the time-varying risk of immune escape in dynamical epidemiological settings.

The phylodynamic curve was originally formulated as a statement on within-host adaptation, encapsulating the non-linear relation between strength of immunity and the risk of a viral escape variant arising during infection. At low levels of immune pressure, viral populations may be large (high abundance) but they face minimal selective pressure and so adaptation is limited. Conversely, as the immune pressure intensifies, the selection for immune escape variants becomes more pronounced, accelerating the rate of viral adaptation. However, beyond a certain threshold, robust immune responses reduce the viral population size enough to restrict the pool of genetic variants and slow adaptation. Consequently, the net adaptation is expected to peak at intermediate levels of strength of immunity. Together, these dynamics produce an inverted U-shaped phylodynamic curve, when the net adaptation rate is plotted against the immune strength.

In this study, we take a population-level view of the phylodynamic curve and consider how the risk of immune escape depends on factors such as population immunity (including factors such as leaky immunity and partial cross-immunity), infection prevalence, transmissibility, rates of case importation etc. The inclusion of imperfect (“leaky”) immunity, a phenomenon that was recently demonstrated in COVID-19 [12], turns out to be especially crucial, as it has profound consequences for the risk of immune escape variant emergence. We provide a quantitative, population-level realization of the phylodynamic curve in stationary situations (such as endemic equilibrium) as well as a framework for computation of the net adaptation rate in arbitrary dynamic settings such as seasonally-driven epidemics, during variant replacement or co-circulation, see Fig. 1 for a schematic. This general and extensible framework allows us to probe the complex dynamics of pathogen-host interactions insofar as they affect the risk of immune evasion.

The findings presented here, and the approach more generally, hold implications for public health, including vaccine design and distribution as well as infectious disease control. Understanding the conditions which lead to heightened risk of escape variant emergence – i.e. the peak of the phylodynamic curve – is crucial. By pinpointing the conditions under which pathogens are most likely to evolve to escape immunity helps prioritize interventions, whether they be in the form of vaccination campaigns, public health policies, non-pharmaceutical interventions or therapeutic approaches.

The phylodynamic curve highlights the heightened potential for variant emergence at intermediate strength of immunity. In accordance with that, the evolutionary consequences of weakly protective immune responses have recently been highlighted by the evolution of SARS-CoV-2, where novel variants tended to represent large genomic jumps relative to the resident variant [13], and a growing body of evidence points to insufficient antibody responses as a key factor in the emergence of these divergent escape variants [14–19].

After introducing the central mathematical model in Section II, we go on to discuss the net adaptation rate in equilibrium in Section III. We begin by considering the simplest scenario, namely the limit of highly durable (albeit imperfectly infection-blocking) immunity. We then consider the more general endemic equilibrium scenario, where immunity duration is finite and protection is imperfect. We then conceptually apply the proposed framework to several different pathogens, characterized by characterized by different degrees of transmissibility, strength of infection and antigenic constraints. A quantitative application of the framework across multiple pathogens may open the door to comparative phylodynamics, improving out understanding of why some pathogens readily evolve immune evading strains, while others do not.

After these equilibrium considerations, we analyze out-of-equilibrium dynamics of the net adaptation rate in Section IV. We consider seasonally driven transmission, and show how importation of cases from less seasonally-varying regions can qualitatively shift the timing of variant emergence. Finally, motivated by the COVID-19 pandemic, we discuss some likely impacts of transmission-reducing non-pharmaceutical interventions (NPIs) on the risk of immune escape variant emergence while the NPIs are in force, as well as after they are lifted.

## II. METHODS

In this section, we derive analytical expressions for the net adaptation rate, defined as the probability per unit time that a viral immune escape variant will emerge. The expressionfor the net adaptation rate depends on the precise underlying epidemiological model and assumptions made, but the derivations follow similar steps in each case.

First, we derive the net adaptation rate in the situation where there is a single resident strain, and examine special cases of this scenario.

Second, we present the net adaptation rate in the presence of two strains with partial cross-immunity, as well as a system of ordinary differential equations describing the dynamics of the strains. In this case, the net adaptation rate describes the probability rate for a *third* (immune-evading) strain emerging.

### A. The net adaptation rate

Assume that a *resident strain* is circulating, with known densities of susceptible (*S*(*t*)), infected (*I*(*t*)) and recovered individuals (*R*(*t*)). Note that these may be time-dependent functions, and are subject to *S*(*t*) + *I*(*t*) + *R*(*t*) = 1. We take immunity to be imperfect (*“leaky”*), meaning that recovered individuals are still partially susceptible, their susceptibility being discounted by a factor of 1 − *σ*. We denote *σ* the *strength of immunity*.

Each infected individual causes new infections at a rate ((1 − *σ*)*R* + *S*)*β*. Immune-evasion mutations are assumed to arise at transmission, with a per-transmission rate of *µ*. Such mutations thus appear in the population at a rate ((1 − *σ*)*R* + *S*)*µβI*. The net adaptation rate measures the rate at which immune-evading mutations appear *and* establish in the population. Thus, we must compute the probability that a newly arisen mutant variant survives (avoids stochastic extinction) or, conversely, the *extinction probability*. To do this, we can use the equation for the extinction probability *d* of a Galton-Watson branching process [20, 21]:

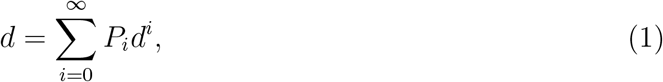

where *P*_*i*_ is the offspring distribution of the mutant variant, and the right-hand side may be recognized as the probability generating function *G*(*d*) associated with *P*_*i*_. The equation for the extinction probability can thus be succinctly written as *d* = *G*(*d*), which is, in general, a transcendental equation.

If transmission is assumed to follow a simple rate process, the offspring distribution will be

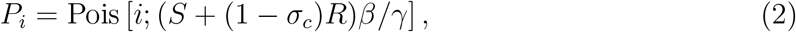

i.e. a Poisson distribution with mean (reproductive number) 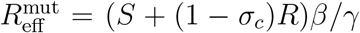, with *γ* the recovery rate and *σ*_*c*_ the strength of cross-immunity between the potential escape variant and the resident strain. This leads to a somewhat opaque expression for the extinction risk, namely *d* = −*W*_0_[−*e*^−*λ*^*λ*]*/λ* where *W*_0_ is the principal branch Lambert *W* function.

However, taking into account that infectious *time* is assumed to follow an exponential distribution in typical compartmental models, the offspring number will follow a *k* = 1 negative binomial distribution (also sometimes referred to as a geometric distribution). In this case, computation of the extinction probability is simplified [22] and yields:

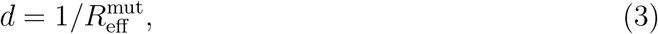

which, in the case discussed above, is simply:

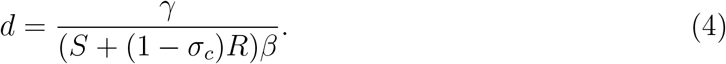

Combining these elements, the net adaptation rate *r*_*a*_ is given by:

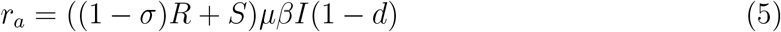

Formally, the extinction risk *d* given above assumes a homogeneous branching process in which the offspring distribution does not change with time. We discuss this simplification in Section A 1 of the Supplement.

Incorporating an influx of mutations from outside the modeled population is straightforward. This amounts to adding a density-independent term to the factor ((1 − *σ*)*R* + *S*)*µβI* which describes the rate at which escape mutations are generated, rendering it ((1 − *σ*)*R* + *S*)*µβI* + *µ*_*im*_. The final net adaptation rate then becomes

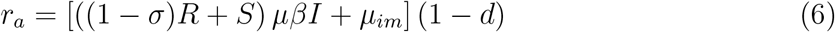

Note that, while *µ*_*im*_ is presently assumed independent of the local densities *S, I* and *R*, it may be modulated by e.g. the incidence at an adjacent location from which the imported cases originate.

The dynamical variables *S, I* and *R* appear in (6) and must be supplied before the net adaptation rate can be computed. Importantly, these fractions may be supplied by empirical timeseries *or* by a dynamical model. In the present work, we focus on model-supplied timeseries, but comment on the prospects of applying the framework to empirical timeseries in the *Concluding remarks* section.

Since the population fractions will be model-derived, we require an epidemiological model with leaky immunity, in line with the assumptions made in the derivation of (6). We will employ a compartmental ODE model including waning, leaky immunity, which we denote SIRS_*σ*_, i.e. a **S**usceptible-**I**nfected-**R**ecovered-**S**usceptible model, extended by including the strength of immunity parameter *σ*. The equations of the SIRS_*σ*_ model are:

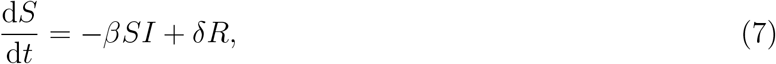

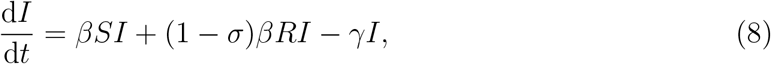

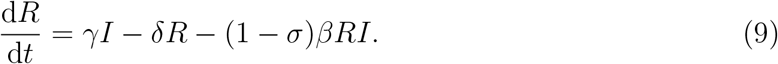

It should be noted that, in our simulations, we will also be interested in scenarios where the transmission rate is seasonally driven. Thus, in addition to the variables *S*(*t*), *I*(*t*) and *R*(*t*), the transmission rate *β* = *β*(*t*) may depend on time as well.

As is clear from the above, the *SIRS*_*σ*_ model yields only the dynamics of the resident strain. This is because we will, in the majority of this paper, only be concerned with the *risk of emergence* of an immune-evading strain on the background of a resident strain. However, in section II B we do extend the framework to evaluate cases where a new strain *does* emerges and circulates at appreciable levels.

a. *Special case: endemic equilibrium*. The endemic equilibrium value of the infected fraction (*I*^∗^) in the SIRS_*σ*_ model can be determined from the equation

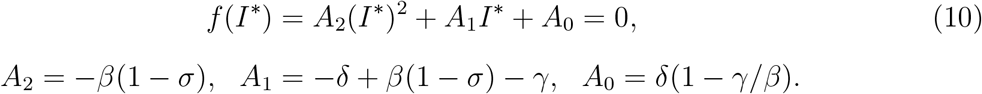 The endemic equilibrium values of the susceptible (*S*^∗^) and recovered (*R*^∗^) fractions can then be determined from

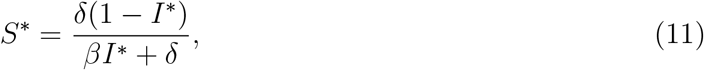

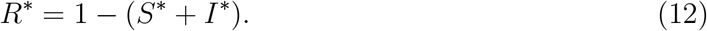 Given the steady-state values *S*^∗^, *I*^∗^ and *R*^∗^, the equilibrium net adaptation rate can thus be computed.
b. *Special case: complete exposure*. A particularly simple special case occurs when the entire population has (at some point) been exposed to the resident strain. This may approximately apply for a disease which produces long-lasting albeit imperfectly infection-blocking immunity. In this case, all individuals are either in compartment *R* or *I*, and the net adaptation rate in the absence of imported cases is given by *r*_*a*_ = (1 − *σ*)(1 − *R*)*Rµβ*(1 − *d*), where we used *R* + *I* = 1. The extinction probability *d* is then derived from a *k* = 1 negative binomial offspring distribution with mean 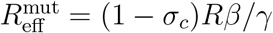.

Since the system is assumed to be at equilibrium, we can solve for the steady-state values of the recovered and infection fractions, *R*^∗^ and *I*^∗^, using

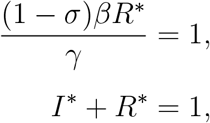

which lead to

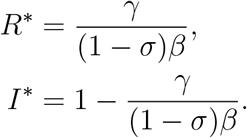

Inserting this in the net adaptation rate yields:

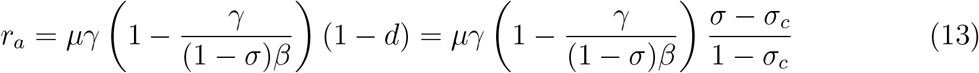

where we used that, in this model

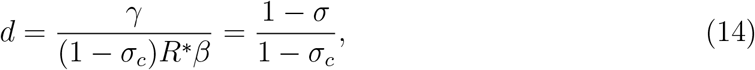

which follows from Eq. (6).

### B. Strain co-circulation

This section tackles the situation where two strains (‘1’ and ‘2’) are circulating at appreciable levels. The resulting model can be used to model the epidemiological dynamics once an immune-evading strain *does* emerge, and to evaluate the risk of another immune evading variant emerging once two variants circulate. In the following, we use *R*_*i*_ (for *i* ∈ *{*1, 2*}*) to refer to individuals recovered from infection with strain *i* only, while the compartment *R* contains individuals who have previously been infected by both types. The infected compartments are of the form *I*_*i*_, corresponding to primary infection with strain *i*, and *I*_*ij*_ (with *i* ≠ *j*), corresponding to infection with strain *i*, having previously been infected with strain *j* (and potentially with *i* itself).

Each infection with strain 1 causes new infections at a rate

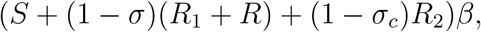

and similarly for strain 2, except with indices 1 and 2 swapped. Mutations due to strain 1 thus arise at a rate

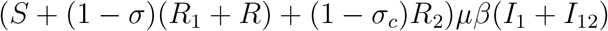

and similarly for strain 2. The net adaptation rate (with a rate of importation, *µ*_*im*_) in the presence of two strains is thus given by:

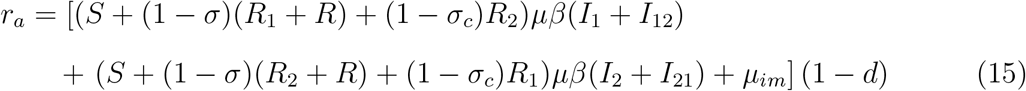

where 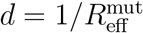 is given by

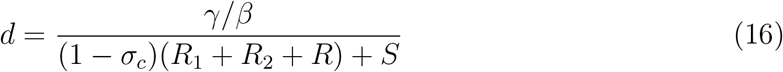

Note that (16) assumes that the potential emerging strain exhibits the same strength of crossimmunity *σ*_*c*_ towards the two circulating strains as these do to each other. This assumption, however, can be easily modified by replacing *σ*_*c*_ in (16) by a distinct cross-immunity strength. Until strain 2 emerges, the net adaptation rate is thus described by (6), and subsequently by (15) (which then describes the per-time risk of a third strain emerging). The system of ordinary differential equations for the two-strain model to which we couple (15) is:

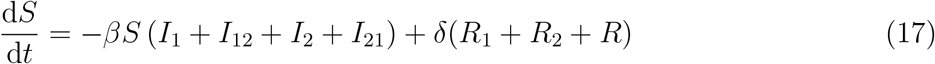

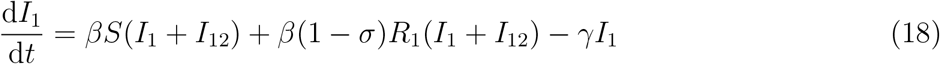

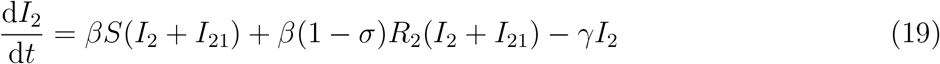

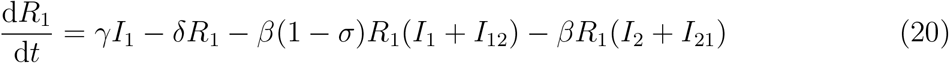

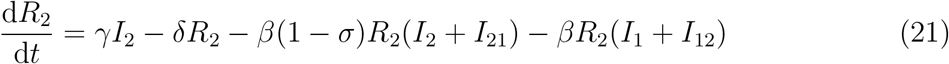

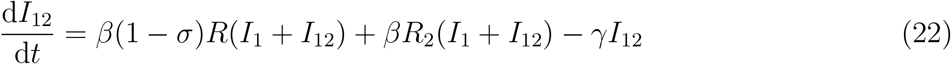

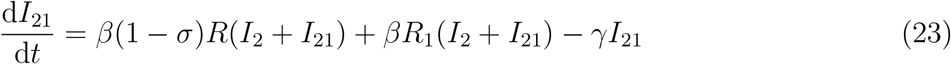

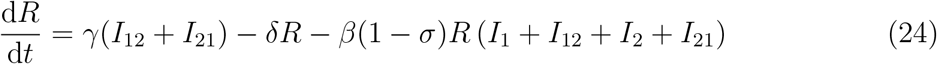

## III. EQUILIBRIUM ADAPTATION RATE

In this section, we explore how the net adaptation rate, i.e. the risk of immune escape, depends the on strength of immunity and transmissibility in steady-state scenarios.

### Complete exposure

We explore two separate equilibrium scenarios of increasing complexity. The first is that of a population that has been completely exposed to the resident variant, meaning that all individuals are either recovered or currently infected. This scenario approximates either the situation after a highly transmissible variant has swept the population, or a disease with long-lasting (albeit imperfect) immunity, to which most individuals beyond a certain age would have been exposed. Just as importantly, this scenario also covers the situation where the population has been vaccinated with a vaccine that does not provide perfect protection against infection [23, 24]. Due to its simplicity, this scenario allows for especially straight-forward interpretation. The resulting family of phylodynamic curves (one for each value of the transmission rate *β*) is shown in Fig. 2A. The inset shows the inverse-U shaped phylodynamic curve that emerges as different values of the strength of immunity *σ* are scanned at fixed transmission rate *β*.

**FIG. 2.**
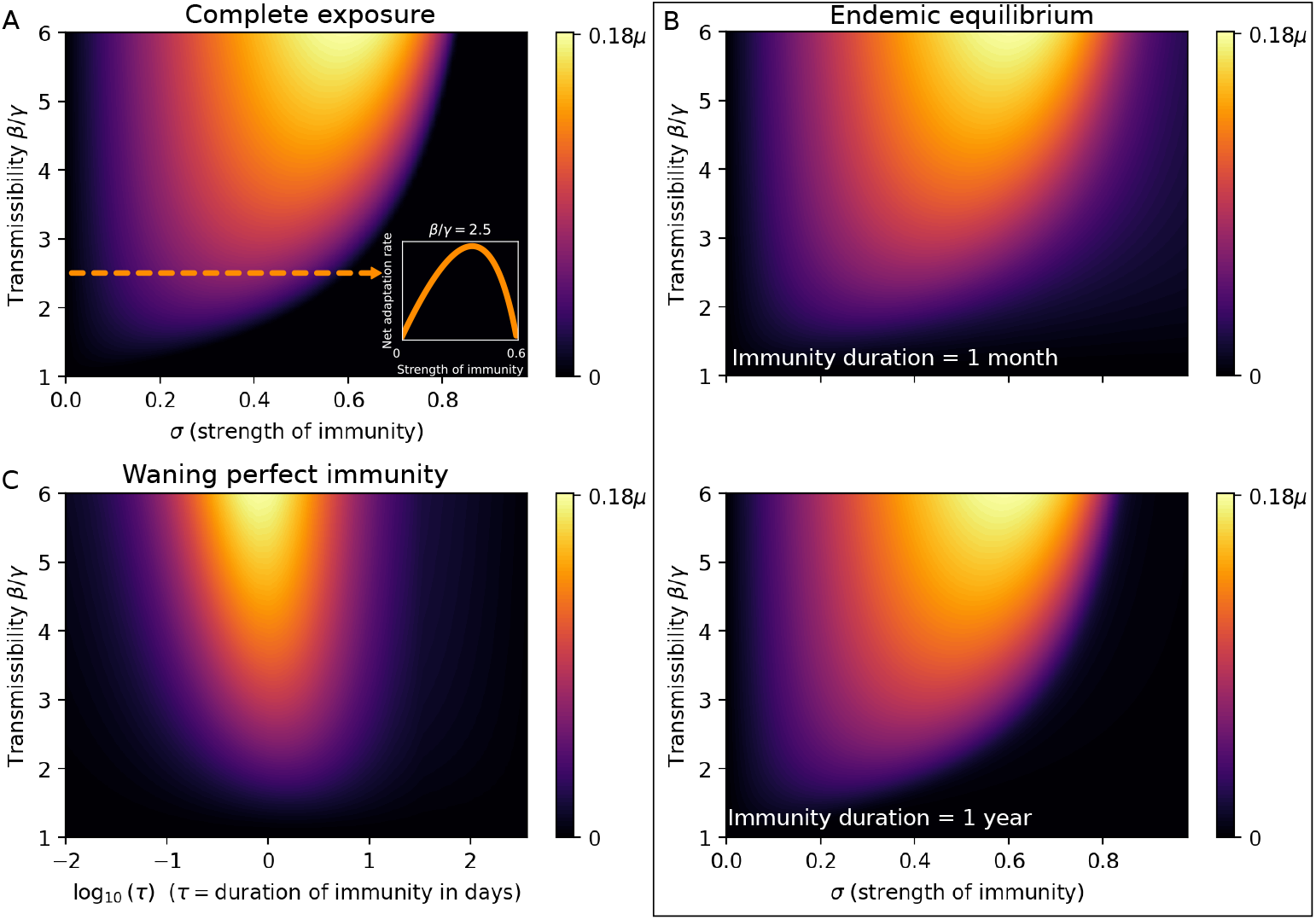
The net adaptation rate in equilibrium scenarios. **A)** The complete exposure scenario, with net adaptation rate given by (13). For each value of the transmission rate *β*, a non-monotonic dependence on the strength of immunity *σ* results. The inset shows the phylodynamic curve that results at *β* = 2.5*γ* corresponding to an *R*_0_ of 2.5 at *σ* = 1. **B)** Endemic equilibrium. The upper panel corresponds to a fast immunity waning rate of δ = 1*/*month, while the lower panel has δ = 1*/*yr. Note that the longer-lasting scenario is quite similar to the complete exposure scenario of panel A. **C)** Waning perfect immunity in endemic equilibrium. The net adaptation rate *r*_*a*_ as a function of transmission rate *β* and duration of immunity, *τ* = 1*/*δ. At fixed *β*, a non-monotonic dependence on δ appears, but maximal *r*_*a*_ occurs at very short immunity duration (approx. a day), meaning that the net adaptation rate is a monotonically increasing function of the waning rate δ at all epidemiologically relevant values of δ, when perfectly infection-blocking immunity (*σ* = 1) is assumed. The recovery rate is fixed at *γ* = 0.5*/*day throughout this figure.

Eq. (13) reveals that, at weak immunity (low *σ*), the net adaptation rate is limited by the mutant survival chance (which simply equals *σ* when cross-immunity is negligible), while at strong immunity (*σ* close to the herd immunity threshold 1 − 1*/R*_0_), the infected fraction is the limiting factor.

In the supplementary material, we numerically simulate the Complete Exposure scenario with explicit stochasticity (i.e. without making any analytical assumptions about the extinction risk, *d*) via the Doob-Gillespie algorithm. In this case, we are able to explicitly measure the net rate of adaptation as the reciprocal of the average time before an immune escape variant emerges. The results are can be found in Fig. S3 and show excellent overall agreement with Fig. 2A.

### Endemic equilibrium

We now turn to the endemic equilibrium solution of the full SIRS_*σ*_ model, which includes waning *im*perfect immunity (i.e. *σ <* 1). Note that the complete exposure scenario previously studied is formally the long-term (equilibrium) solution to the SIRS_*σ*_ model in the limit where δ tends to 0. Fig 2B shows the net adaptation rate (as a function of *β* and *σ*) at two different rates of waning, corresponding to mean durations 1*/*δ = 1 month and 1*/*δ = 1 yr respectively. The recovery time is 1*/γ* = 5 days, and we note that already at an immunity duration of 1 month, the plot bears a strong resemblance to the complete exposure result of Fig. 2A, and the two are almost indistinguishable at an immunity duration of 1 year.

### Evolutionary consequences of waning immunity vs. leaky immunity

In the previous sections, we have assumed that immunity is imperfect, in the sense that it reduces susceptibility by a factor (1 − *σ*) rather than completely. This is in contrast to most theoretical models of how the evolution of immune escape is shaped by population immunity, which have typically assumed that homologous immunity is perfect while it lasts [25–27], and that variable “strength” of immunity can be captured by varying the *duration* of protection. In this section, we will adopt the assumption that homologous immunity is perfectly infection-blocking while it lasts, and is thus limited only by exponential waning. In this case, immunity is then binary: either one is perfectly protected against infection, or one is fully susceptible [28]. We explore this assumption to see how it affects conclusions regarding the net adaptation, and thus the evolution of immune escape.

Assuming endemic equilibrium, we explore the effects of waning immunity in Fig. 2C – note the logarithmic horizontal axis. While a non-monotonic relation between rate of waning and net adaptation rate is observed, the downslope only appears at waning rates in excess of 100*/*yr which are epidemiologically irrelevant. We thus conclude that exponentially waning perfect immunity does not yield an inverse U-shaped phylodynamic curve, and that within epidemiologically relevant parameters, faster waning tends to lead to faster adaptation. This finding may lend itself to at least crude empirical testing, by comparing pathogens with different rates of immune waning. However, it should be mentioned that this result is likely to depend, to some degree, on assumptions made about the per-generation rate of antigenic evolution vs. the per-infected-time rate – an aspect that warrants exploration in future works. Overall, we are led to conclude that the distinction between leaky immunity and waning but polarized (all-or-nothing [28]) immunity has profound consequences for the emergence of escape variants, highlighting the need for mapping the individual-level post-exposure (and post-vaccination) protection against reinfection for circulating pathogens. While leaky immunity has been demonstrated in COVID-19 [12], a challenge study with infectious hematopoietic necrosis virus in rainbow trout found very heterogeneous vaccinal immunity, in an almost all-or-nothing fashion [29], emphasizing the multiple ways in which protection may be imperfect.

### The role of antigenic constraint and the effective mutation rate

Due to the simplicity of the proposed framework, molecular aspects of antigenic evolution are not explicitly modeled and as a consequence, the within-host evolutionary process is highly coarse-grained. However, through the effective mutation rate *µ*, which parameterizes the per-transmission risk of an immune-evading mutation occurring, we can probe some of the overall effects of antigenic constraint.

Despite facing strong selective pressure and maintaining substantial viral abundance, some viruses exhibit limited antigenic variability [30, 31]. This situation is well-exemplified by measles. With its high basic reproductive number, strong, durable immunity and relatively widespread vaccination, measles would be expected to present ideal conditions for the emergence of escape variants. However, such variants are not observed. This phenomenon can largely be attributed to “antigenic constraint”, where biophysical limitations restrict the virus’s capacity for viable mutations. In the case of measles, these constraints might arise from conformational limitations of its primary antigens, where substantial alterations could impede the virus’s infectivity or replication efficiency. Despite observed genetic variability in the F (*fusion*) and H (*hemagglutinin*) envelope proteins of measles, analysis has shown that existing constraints only allow for a single serotype [32]. The consequences of these constraints are remarkable, exemplified by the fact that a genotype capable of evading vaccine-induced immunity has not emerged after decades of widespread vaccination [33]. Serum neutralization assays with mutated H proteins has shown that several co-dominant glycoprotein epitopes exist within this protein, and that the H protein is co-dominant with respect to serum neutralization with the F protein, meaning that a large number of mutations (each with potential fitness costs) would be required to effectively evade immunity [33, 34].

Similarly, influenza A/H3N2’s antigenic evolution is not without its bounds. The evolution of its hemagglutinin has been shown to be shaped by geometric constraints [35]. And as detailed in [36], constraints on net charge – maintained by the balancing of charged amino acids – limit the evolution of the neuraminidase antigenic region studied. A similar result has been found the hemagglutinin of influenza A/H1N1 [37].

Together, these examples illustrate an important aspect of pathogen evolution: the balance between selective pressures and the inherent structural and functional constraints of the pathogen. In our model, we only very crudely incorporate antigenic constraints, and do so via an effective description: we subsume their effects into the *µ* parameter, the antigenic mutation rate. We explore the effects of this parameter in Fig. 3, where we have conceptually placed a number of pathogens in a multidimensional net adaptation rate landscape, parameterized by the effective mutation rate *µ*, the transmission rate *β* and the strength of immunity *σ*. We stress that this rough overview is only meant to show how the proposed framework can be used to create an easily interpretable immune escape risk classification.

**FIG. 3.**
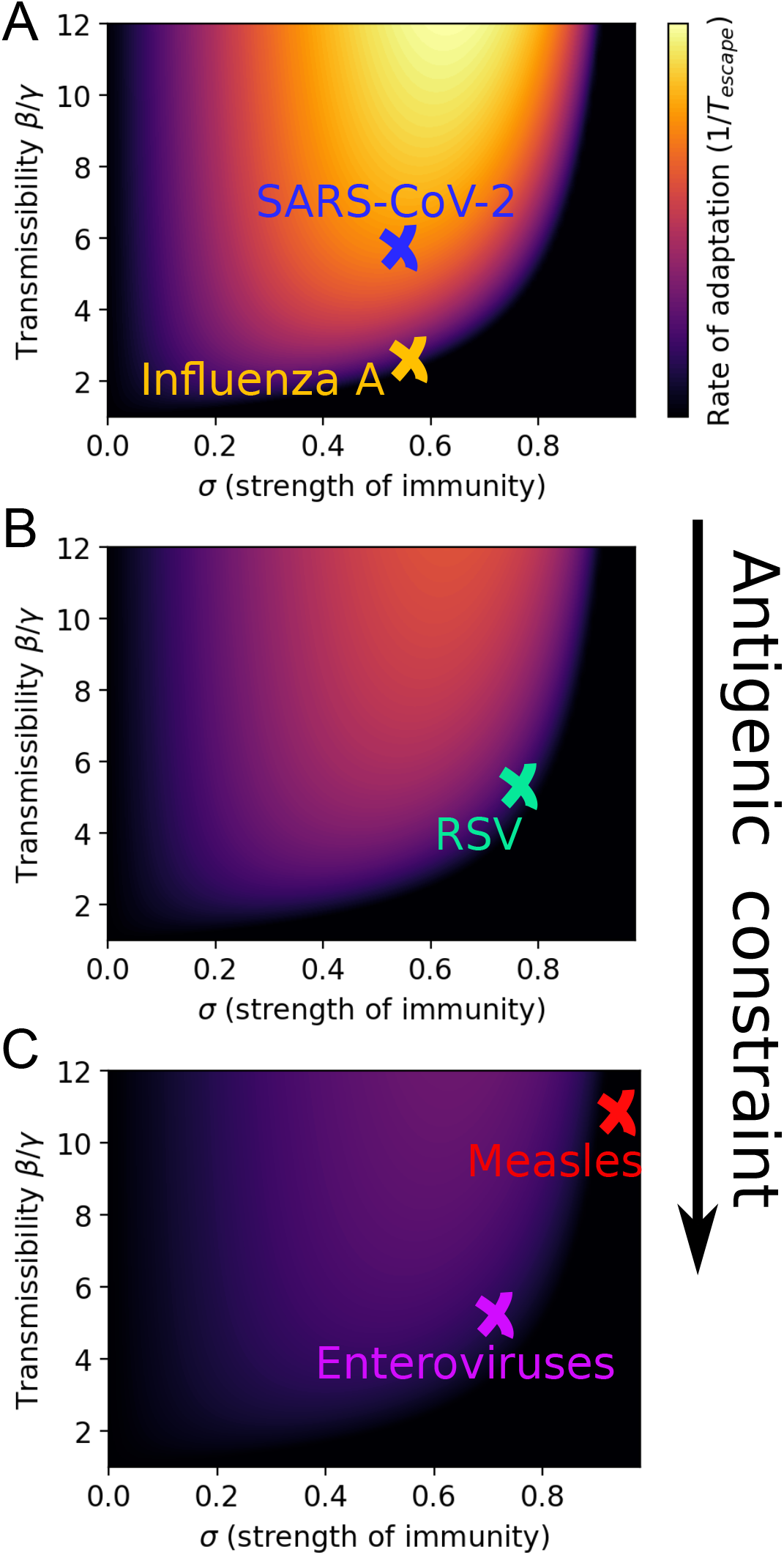
A conceptual phylodynamic landscape, parameterized by transmissibility *β*, strength of immunity *σ* and effective mutation rate *µ*. Stronger antigenic constraints decrease the effective mutation rate, as indicated by the vertical arrow. **A)** *µ* = 1, **B)** *µ* = 0.6, **C)** *µ* = 0.3. The recovery rate is fixed at *γ* = 0.5*/*day throughout this figure. In this figure, no cross-immunity between the resident variant and a putative escape variant is assumed. Fig. S2 explores the impacts of partial *cross*-immunity.

The basic reproduction number for influenza (*R*_0_ ≈ 2) was based on [38]. *R*_0_ of course refers to the reproductive number in a naive population. For influenza, this is best approximated by measurements of pandemic influenza while seasonal influenza typically fails to realize this figure, precisely because of partial immunity. For enteroviruses, a very diverse class, estimates vary widely. [39] obtain estimates for EV-D68 in the 10-20 range, while [40] estimate the values 5.5 for EV-A71 and 2.5 for Cox A16. [41] report an *R*_0_ of 5.0 for Cox A6, 2.4 for Cox A16: 2.4 and 3.5 for EV-A71. For respiratory syncytial virus (RSV), estimates are also highly variable. [42] finds a mean literature estimate of approx. 4.5 while they themselves estimate it to 2.8. [43] estimates *R*_0_ at 3.0, while [44] find a higher value of 8.9. SARS-CoV-2 *R*_0_ values vary by variant, with the ancestral strain estimated at around 3 [45], while the *R*_0_ of the Omicron variant has been estimated at 8-9 [46].

## IV. NON-EQUILIBRIUM DYNAMICS

In this section, we allow full temporal variation in the susceptible, infected and recovered population fractions, and allow for time-dependent transmission rate *β*(*t*).

### A. Seasonality, imported cases and variant emergence

Seasonal variation in transmission rate not only leads to periodic outbreaks, but to a strongly time-varying net adaptation rate as well. Fig. 4 shows the net adaptation rate as given by (6) coupled to the system of differential equations (7)–(9) describing the SIRS_*σ*_ model, with sinusoidally varying transmission rate *β*(*t*) = *β*_0_(1 + 0.2 cos(2*πt/*365d)). In panel A, the importation rate is turned off (*µ*_im_ = 0), a scenario which leads to the bulk of the emergence risk occurring during the growth phase of the seasonal epidemic (note that the risk of emergence is very low during the decreasing phase of the epidemic, echoing a result by [47]). This is in contrast to the case in panel B, where a constant importation rate is turned on (*µ*_im_ = 0.025*µ*) and a large part of the emergence risk (understood as the area under the red curve) occurs during the *trough period*, i.e. while prevalence is locally practically non-existent. This shows one of the potentially profound consequences of year-round reservoirs for otherwise seasonal pathogens. A salient example may be influenza A, which is strongly seasonal in temperate climates, while tropical regions exhibit extensive year-round influenza activity as well as irregular timing of outbreaks [48–51].

**FIG. 4.**
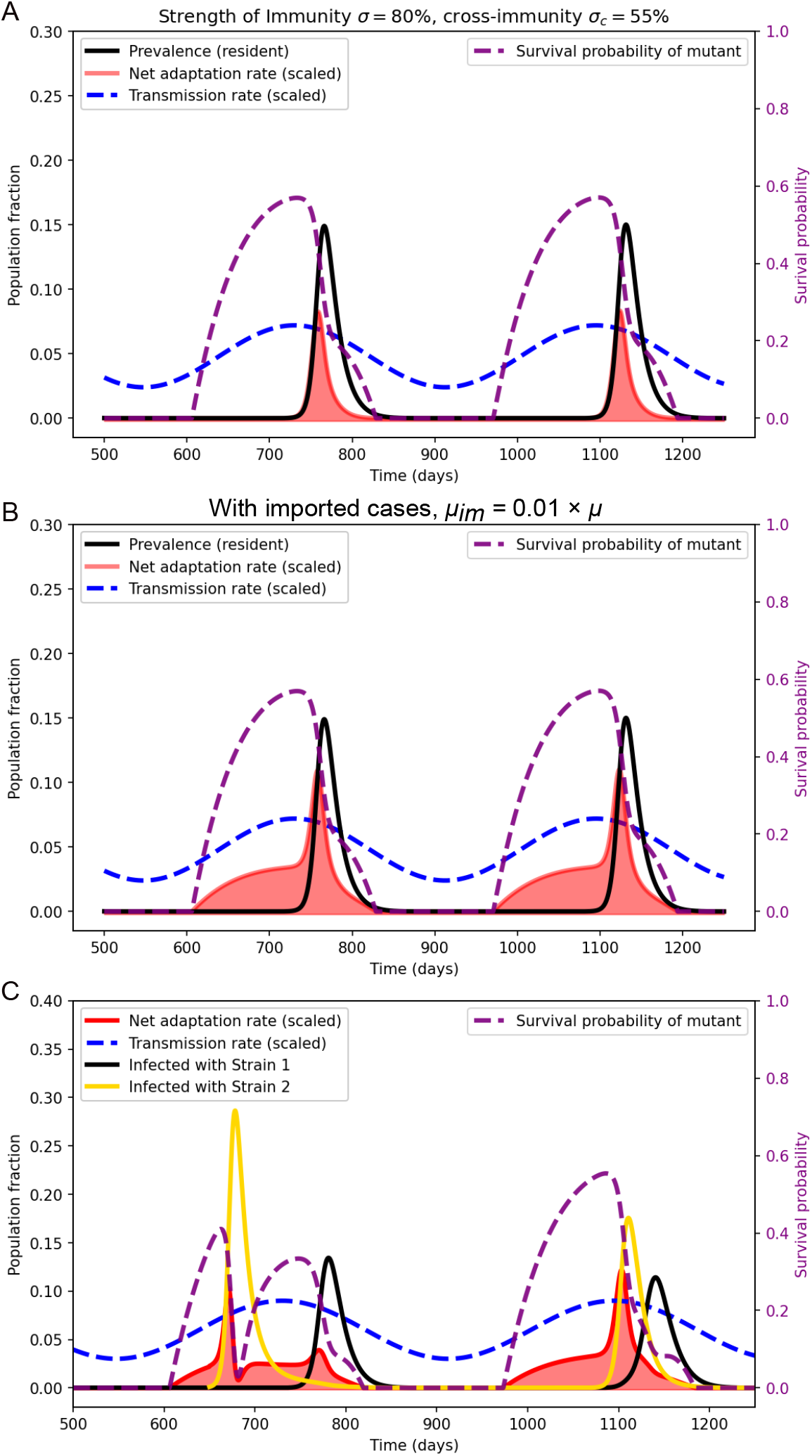
Seasonality and the emergence of an escape variant. Sinusoidally modulated transmission rate (dashed blue curve). **(A)** No imported cases (*µ*_im_ = 0). The bulk of the emergence risk (area under the red curve) occurs during the epidemic growth phase, while incidence is high and the susceptible fraction is substantial. **(B)** Constant importation rate *µ*_im_ = 0.01*µ*, modeling imports from a region with year-round circulation. The trough period now sees a substantial emergence risk. **C)** Full two-strain model. An immune-escape variant (**yellow** prevalence curve) emerges at time *t* = 650 days and disrupts the seasonal pattern. The moderate cross-protection (*σ*_*c*_ = 0.55) leads to subsequent co-circulation.

In Fig. 4C, the two-strain model described by equations (17)-(24) is implemented, and an immune escape variant (“Strain 2”) emerges at time *t* = 650 days. The surge of the new strain disrupts the seasonal pattern and the moderate level of (symmetric) cross-protection (*σ*_*c*_ = 0.55) allows for subsequent co-circulation of the newly emerged and the previously dominant strain (“Strain 1”). It is not just the pattern of circulation that is disrupted – so is the net adaptation rate, exhibiting two peaks within a single season. The simulation of Fig. 4C also highlights a central aspect of the proposed model framework: the net adaptation rate describes the potential for adaptive sweeps, but once such a sweep *occurs*, it will strongly perturb the forward dynamics.

### B. The impact of non-pharmaceutical interventions

The non-pharmaceutical interventions (NPIs), such as lockdowns, implemented during the COVID-19 pandemic were often highly effective in reducing the COVID-19 transmission rate [52], as well as those of other circulating respiratory pathogens [53–59]. While the short-term epidemiological implications for COVID-19 are reasonably well-understood, the effects on pathogen evolution are likely complex and remain relatively poorly understood, especially as it pertains to pathogens other than COVID-19, whose dynamics were also affected by interventions.

In this section, we first explore how NPIs acutely affect the net adaptation rate in a few simple situations.

In Fig. 5, we assume that 90% of the population are initially vaccinated with a moderately effective vaccine that reduces susceptibility by 80% against the resident strain and by 55% against a potentially emergent strain. We also assume that immunity (vaccine-induced as well as infection-derived) wanes at a rate of 1yr^−1^. We include the initial vaccination to show the effects of immune waning during periods where no (or very little) circulation is taking place locally. At *t* = 50days, NPIs are introduced either globally (panel A) or only locally (panel B), lowering the transmission rate *β* (blue dashed curve in the figure). In the former case, NPIs are able to keep the net adaptation rate at essentially zero, since the implemented interventions effectively limit the viral abundance. In the latter case, NPIs initially hamper adaptation, but once immunity wanes even moderately, the net adaptation rate begins to increase. This is due to the favorable growth conditions created for an invading escape variant – no major competition from a resident variant, but sufficient numbers of fully or partially susceptible individuals for a somewhat immune-evading variant to invade. The importation rate from neighboring (intervention-free) regions then continuously provide the “sparks” which can ignite such an invasion. In this sense, imported cases effectively bypass the low local viral abundance that would otherwise limit the net adaptation rate [60].

**FIG. 5.**
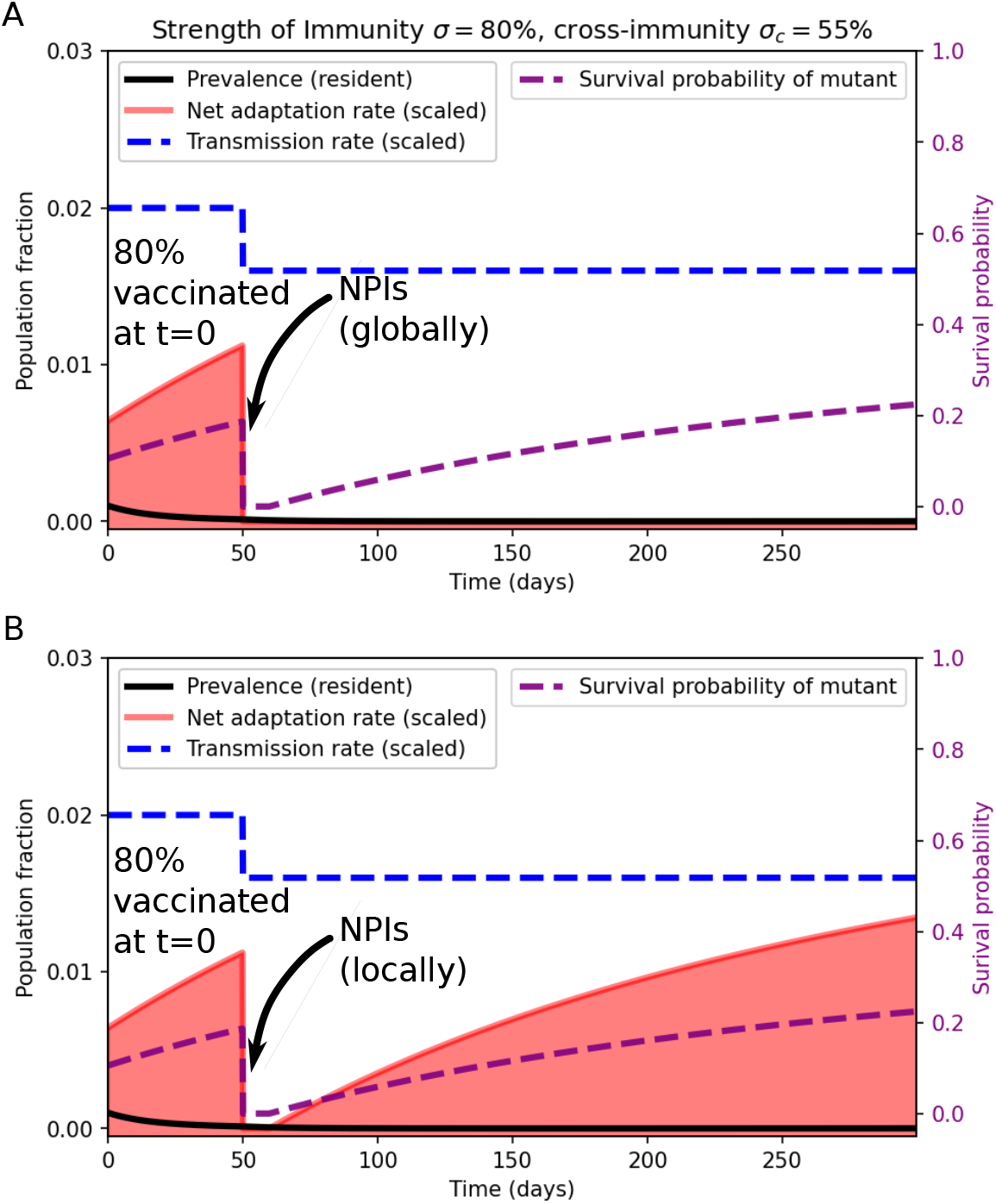
Impact of non-pharmaceutical interventions on the net adaptation rate (red curve), when applied globally **(A)** and locally **(B)**. In the local case, a constant importation rate of *µ*_*im*_ = 0.01*µ* is assumed.

Having considered the acute effects of non-pharmaceutical interventions, i.e. the effects they have while they are in force, we move on to the question of what happens once NPIs are lifted and the transmission rate increases again. We will specifically consider the effects on non-COVID-19 pathogens, which were also affected by NPIs but were not as extensively vaccinated against during the NPI period. The lifting of restrictions has led to surges in several pathogens [61] due to the build-up of susceptibles during the NPI period. We find (Fig. 6A) that such a surge is accompanied by a rapidly rising net adaptation rate, despite the initially low immune pressure.

**FIG. 6.**
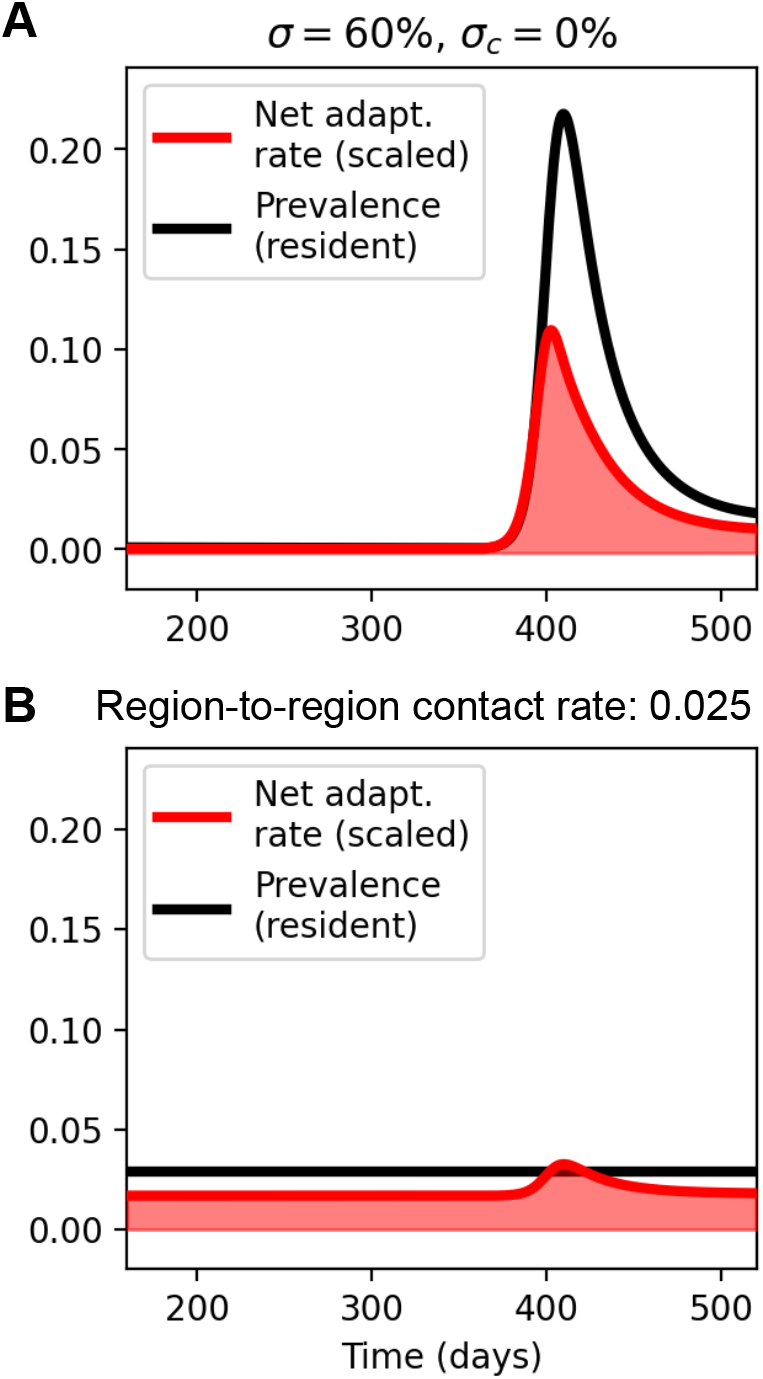
Net adaptation rate dynamics (of non-COVID respiratory pathogen) once non-pharmaceutical interventions are lifted, in the region previously under NPIs (the *Post-NPI* region)**(A)** and in an adjacent region not previously subjected to extensive restrictions (the *Never-NPI* region) **(B)**. The importation rate in the Never-NPI Region is assumed proportional to the prevalence in the Post-NPI region: *µ*_im_(*t*) = *r*_*c*_*µI*_Post-NPI_(*t*), with *r*_*c*_ = 0.025 a contact rate from the Post-NPI to the Never-NPI region.

However, viewing a previously locked-down region in isolation hides the effects that such a local surge may have on neighboring regions with different levels of population immunity. Specifically, regions which did not experience extensive lockdowns may have appreciable population immunity towards the pathogen in question, and correspondingly different selection pressures. To model this, we consider a region that never implemented NPIs (the *Never-NPI Region*), and in which the disease in question is instead in an endemic equilibrium. During the surge in the previously locked-down region (the *Post-NPI Region*), some spill-over to neighbouring regions is likely. Thus, we model an importation rate in the Never-NPI Region that is proportional to the prevalence in the Post-NPI region: *µ*_im_(*t*) = *r*_*c*_*µI*_Post-NPI_(*t*), with *r*_*c*_ a contact rate from the Post-NPI to the Never-NPI region. We observe that, although the prevalence in the Never-NPI region stays constant, the region-to-region contact leads to an appreciably elevated net adaptation rate in the Never-NPI region as well (Fig. 6B).

The impact of cross-immunity (infection-derived as well as vaccine-induced) that reduces susceptibility (and infection duration) on the growth rate of an invading variant and the eventual depth of the trough it undergoes was considered by [62]. However, the authors do not consider imperfect *homologous* immunity, which we have found to be an important factor in shaping the risk of immune escape. It is worth noting that they find that the depth of the trough that an invading variant undergoes follows a U-shaped curve as a function of strength of immunity (as opposed to the *inverse* U-shaped phylodynamic curve). This is expected: highly successful variants tend to overexploit the “resources” (susceptibles), and then undergo a severe trough. However, in practice, deep local troughs are typically not enough to eradicate a novel escape variant, since out-of-phase outbreaks will typically occur elsewhere, negating the effect of the trough. The exception to this rule being the implementation of global or almost-global NPIs, which would synchronize the troughs, similar to what we considered in Fig. 5. If such troughs are sufficiently deep and synchronous, previously widespread pathogen lineages may die out. An example of this may have been observed, with the probable extinction of the Yamagata lineage of Influenza B virus [63].

## V. LIMITATIONS

Our transmission model assumes mean-field behavior, relying on a homogenous mixing assumption and neglects any person-to-person heterogeneities in e.g. susceptibility and infectiousness. While these are often reasonably accurate assumptions [64, 65], it has been shown that departures from uniform infectiousness can have profound implications for epidemic dynamics, especially when social contact networks are restricted (such as during lockdowns), representing a departure from homogeneous mixing [66–68]. Evolutionary models have suggested that the two may even be coupled by evolutionary pressures, with NPIs affecting the evolutionarily favored transmission patterns of pathogens [69].

While we have included both imperfectly infection-blocking and finite-duration (“waning”) immunity in the model presented here, a more detailed model of partial immunity is likely to lead to further insights. In [70], the authors use an antibody titer model to describe partial immunity, a feature that could possibly be included in a more elaborate model of the net adaptation rate.

Diversity in vaccine-induced immune responses constitutes another source of heterogeneity that we did not include, which may affect the establishment probability of escape variants. This phenomenon was studied by [71], who introduced the concept of *escape depth* to describe the fraction of escape mutants likely to spread in a vaccinated population.

Finally, and perhaps most importantly, our highly idealized model does not include any detailed antigenic fitness landscape. Studies on the relation between antigenic and genomic distance suggest a correlation between the two [72, 73], while antigenic cartography indicates that variants are typically well-separated in antigenic space [74–77]. Both of these findings challenge the memory-less constant-rate process assumption we make in this paper. We anticipate that it may be possible to refine the model by more explicitly modeling genomic and antigenic distances and incorporating data on typical relations between the two.

## VI. CONCLUDING REMARKS

In this study, we have proposed a framework to explore the dynamics of viral escape variant emergence, in which a population-scale phylodynamic curve emerges dynamically, encapsulating the non-linear relationship between population immunity and viral adaptation. The framework contributes to a broadening understanding of the conditions under which viral immune escape variants are most likely to emerge, providing valuable insights for public health interventions and vaccine strategies.

We have shown that, far from being a static quantity, the same mathematical expression of the net rate of adaptation that gives rise to the phylodynamic curve can straight-forwardly be coupled to dynamical transmission models. This then allows evaluation of the risk of viral escape variant emergence in highly dynamical settings, including during public health interventions and seasonal variation.

We stress that the expression given for the net adaptation rate can be applied to *empirical timeseries*. The equation involves the population fractions of susceptible, infected and recovered individuals, and while we have used dynamical models to supply these quantities, nothing prevents them from being supplied by empirical observations. This, however, does hinge on relatively high-fidelity suceptible reconstruction being performed, including the effects multiple variant exposures and vaccinations where relevant. To effectively relate our model framework to real-world pathogens, several types of data will thus be essential. Accurate data on recovery rates, and transmission rates and infection prevalence are of course crucial for applying the model in any given scenario. Perhaps more crucially, the model relies on good estimates of the strength and duration of (cross-)immunity and the ability to perform (multi-variant) susceptible reconstruction, and thus benefits from input from serological studies.

The framework we propose offers a foundation for emergence risk evaluation. By applying the model across various pathogens, we may gain insights into the differing propensities of viruses to develop immune escape variants. This comparative approach is likely to elucidate the factors that make certain pathogens more susceptible to rapid evolution, thereby informing targeted surveillance efforts and enhancing intervention strategies. Moreover, our model can serve as a tool for optimizing vaccine design and distribution. By simulating different vaccination strategies, the framework may help identify the conditions that minimize the net adaptation rate.

Finally, the evolutionary consequences of the COVID-19 non-pharmaceutical interventions across circulating pathogens are still being examined and evaluated. Our proposed framework can form part of a theoretical basis for these evaluations, providing insights into how interventions may influence the evolutionary dynamics of multiple pathogens.

## ACKNOWLEDGEMENTS

We have become aware that James Bull, Katia Koelle and Rustom Antia have independently developed a distinct approach to addressing questions related to the emergence of immune escape variants. Their forthcoming work, *“Waning immunity drives respiratory virus evolution and reinfection”* [78], addresses a set of partially overlapping problems, albeit in a very different framework. After consultation, we have agreed to coordinate the submission of our respective manuscripts to bioRxiv.

BFN acknowledges financial support from the Carlsberg Foundation (grant no. CF23-0173). CMSR acknowledges funding from the Miller Institute for Basic Research in Science of UC Berkeley via a Miller Research Fellowship. BTG acknowledges support from the Princeton Catalysis Initiative and Princeton Precision Health. We thank the High Meadows Environmental Institute (HMEI) at Princeton University for support.

## Appendix A: Supplementary Material

### 1. Extinction risk and the quasi-stationarity criterion

As remarked above, the net adaptation rate depends on the *extinction risk d*, which we compute using a result from the theory of branching processes [20, 21], namely that the infinite-time extinction risk for a branching process with a time-invariant offspring distribution *P*_*i*_ satisfies the equation

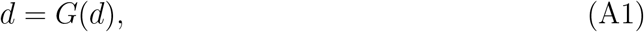

where *G*(*d*) is the probability generating function associated with the probability function *P*_*i*_.

To see how this comes about, let us denote the number of offspring in the *n*’th generation by *i*_*n*_. The probability for extinction in *n* generations (or less) is thus the quantity *d*_*n*_ = Pr[*i*_*n*_ = 0], where Pr[*·*] denotes a probability. Similarly, the infinite-time extinction risk is given by

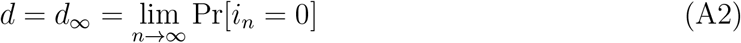

To compute *d*_*n*_, observe that the possible ways to achieve extinction no later than the *n*’th generation are:

- Having 0 offspring (immediate extinction), which happens with probability *P*_0_,
- having one offspring whose lineage dies out in (no more than) *n* − 1 generations, which happens with probability *P*_1_*d*_*n*−1_,
- having two offspring whose lineages both (independently) die out in (no more than) *n* − 1 generations, which happens with probability 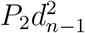,
- and so on.

Adding up these probabilities, we find the expression

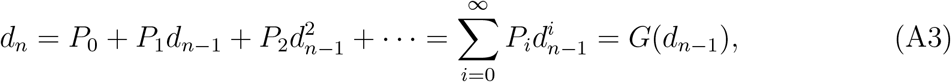

where, in the last step, we recognized the probability generating function associated with *P*_*i*_. From this, the infinite-time result *d* = *G*(*d*) also directly follows, since both *d*_*n*_ and *d*_*n*−1_ tend to *d*_∞_ = *d* as *n* tends to ∞. Note that the equation *d* = *G*(*d*) is, in general, a transcendental equation, which may be solved either numerically or graphically.

In using the parameter *d* in (6), we are assuming a kind of quasi-stationarity, namely that the offspring distribution *P*_*i*_ changes slowly enough that we can consider it constant during the time required for extinction/establishment of a novel variant. The time required may be formally infinite (since we have let *n* → ∞). However, we can test the robustness of this assumption by considering the value of *d*_*n*_ for moderate values of *n* and see how well it approximates *d*. In order to do this, we note that *d*_*n*_ = *G*(*d*_*n*−1_) is a first-order recurrence relation, and that a boundary condition is given by *d*_0_ = *P*_0_, i.e. the probability of going extinct immediately by having no offspring. From this, the subsequent values of *d*_*n*_ for *n* ≥ 1 may be determined iteratively.

In Fig. S1, the extinction probability after *n* generations, *d*_*n*_, is shown for a range of average reproductive numbers (avg. number of offspring). The plots show that the difference between *d*_*n*_ and *d* = *d*_∞_ is quite low as long as *n* is greater than 3 or 4. This has implications for the quasi-stationarity requirement mentioned above, since it shows that the survival probability 1 − *d* is approximately realized after just a few generations. Thus, we do not require that the offspring distribution remains constant, but only that it does not change drastically within the first few generations after the initial emergence of the escape variant in a single infected individual. Note that the extinction probability *d*_*n*_ takes the longest to stabilize when the mean reproductive number is around 1.

### 2. Stochastic (Doob-Gillespie) simulation of the emergence process

In this section we simulate the stochastic emergence of an immune escape variant using the Doob-Gillespie algorithm [79], without assuming eq. (6). We will be focusing on the Complete Exposure scenario (as described in the Methods section). The Gillespie system is described by the following rate equations:

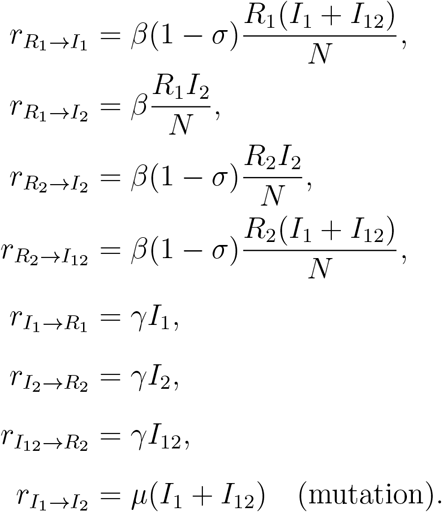

The system is a two-strain system similar the one described in the Methods (Eqs. (17)-(24)), but a few simplifications have been made: since we are considering the Complete Exposure scenario, no fully susceptible class is included. Similarly, we are only interested in whether a potential immune escape variant (strain 2) establishes, but not in the dynamics once it has infected a sizable fraction of the population. Note that we have assumed no cross immunity here, *σ*_*c*_ = 0.

Since we assume complete exposure to strain 1 initially, we include the *I*_12_ compartment, which includes those infected with strain 1 after infection with strain 2, but do not explicitly include *I*_21_, since all individuals are assumed to have been exposed to strain 1 anyway (and thus this class is subsumed in *I*_2_).

In order to assess emergence risk, we record the time *T*_escape_ until an immune escape variant emerges in each simulation, and the net rate of adaption given is then the average value of 1*/T*_escape_ across 1000 simulations for a given parameter value. We consider a variant to have “emerged” (i.e. appeared and avoided stochastic extinction) once it reaches a prevalence of 100 individuals. The resulting heatmap can be seen in Fig. S3. We note that it is qualitatively very similar to what we obtain using the analytical framework (e.g. Fig. 2A).

**FIG. S1.**
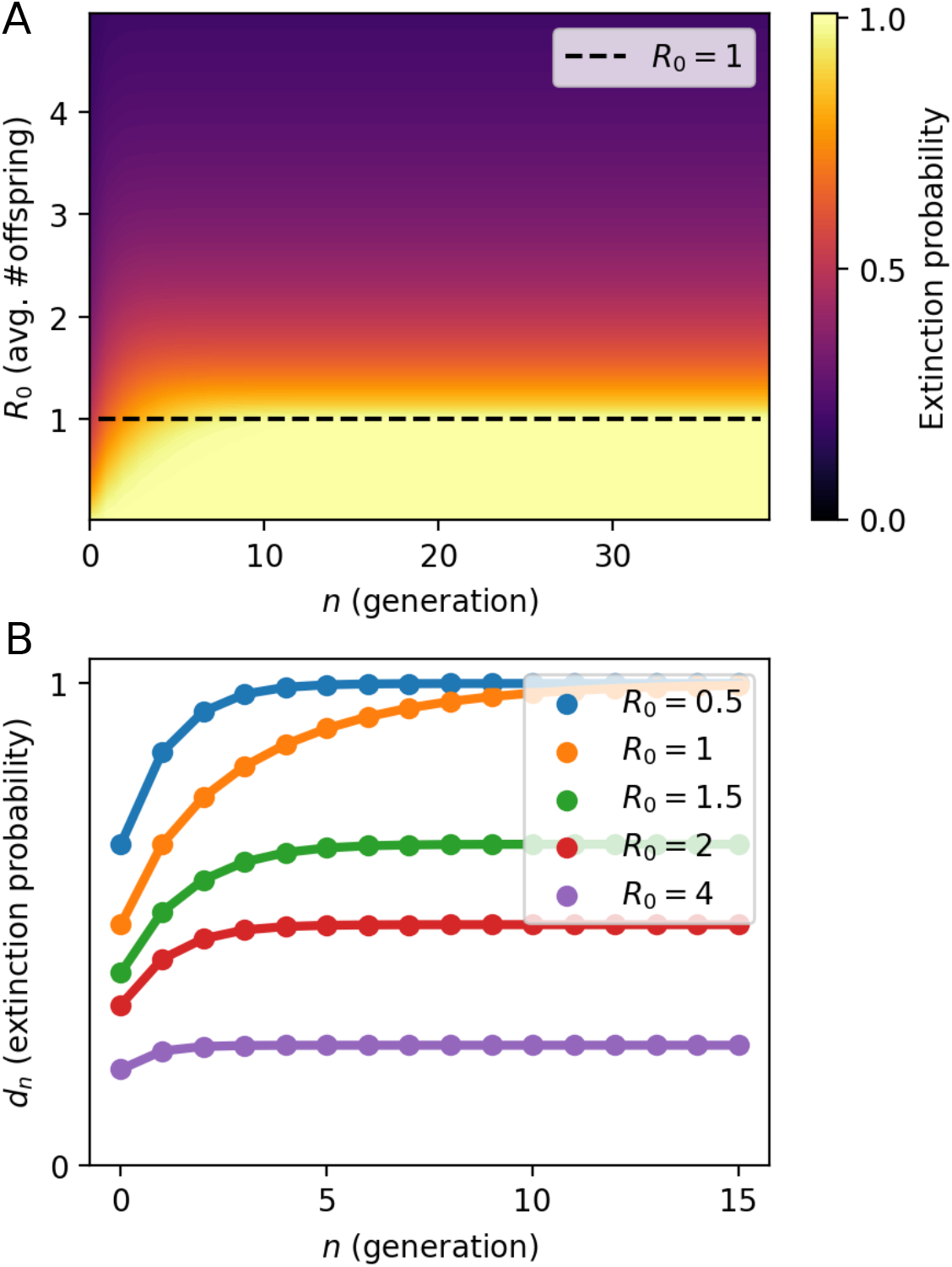
Extinction Probability of a branching process with geometric offspring distribution after *n* generations, as a function of *R*_0_. **A)** The extinction probability, across many values of the mean reproductive number (*R*_0_) and number of generations. The color indicates the probability of the branching process going extinct in *n* generations or less. **B)** The extinction probability *d*_*n*_ as a function of generation *n*, for a few select *R*_0_ values. Beyond the first few generations, the extinction probability changes little.

**FIG. S2.**
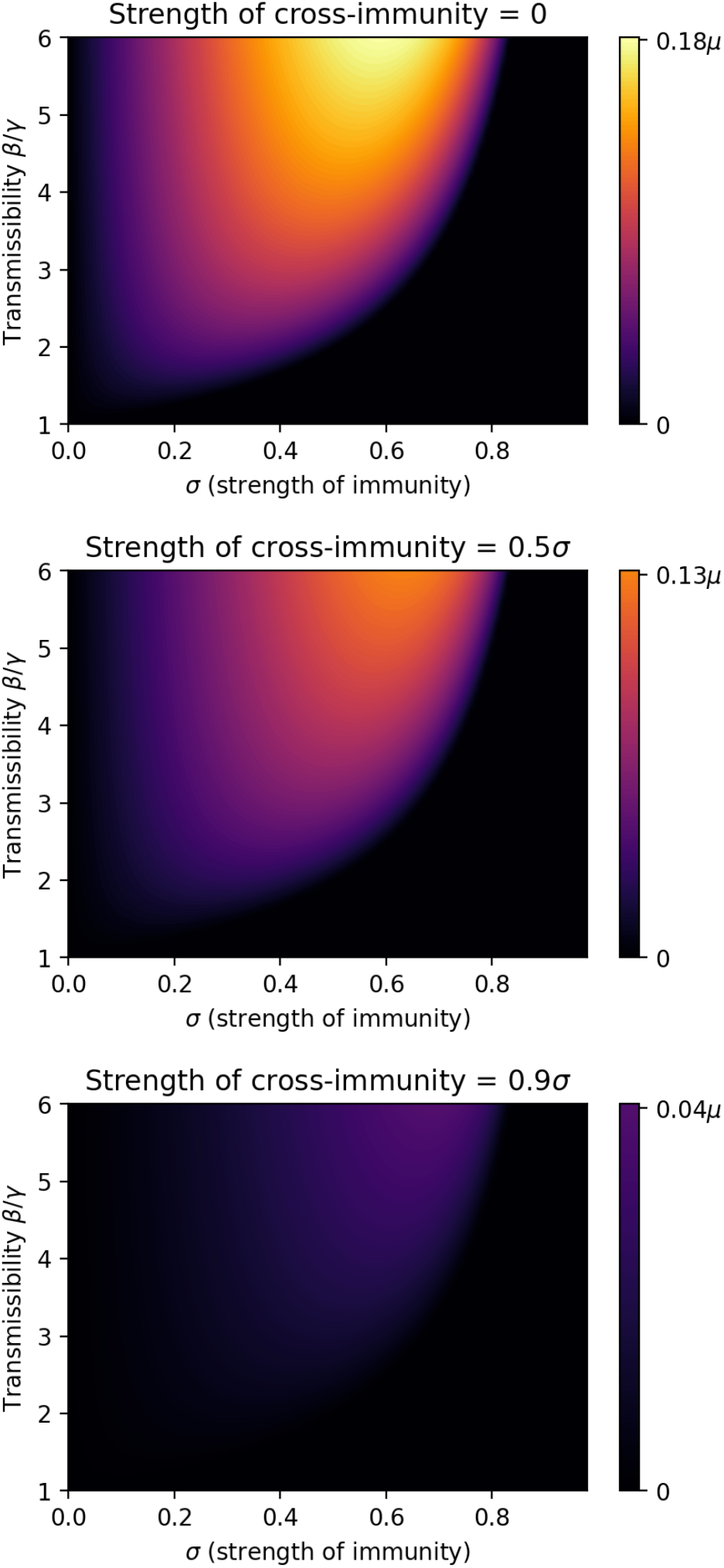
The impact of partial cross-immunity. *σ*_*c*_ (heterologous immunity) on the net adaptation rate. 0.6

**FIG. S3.**
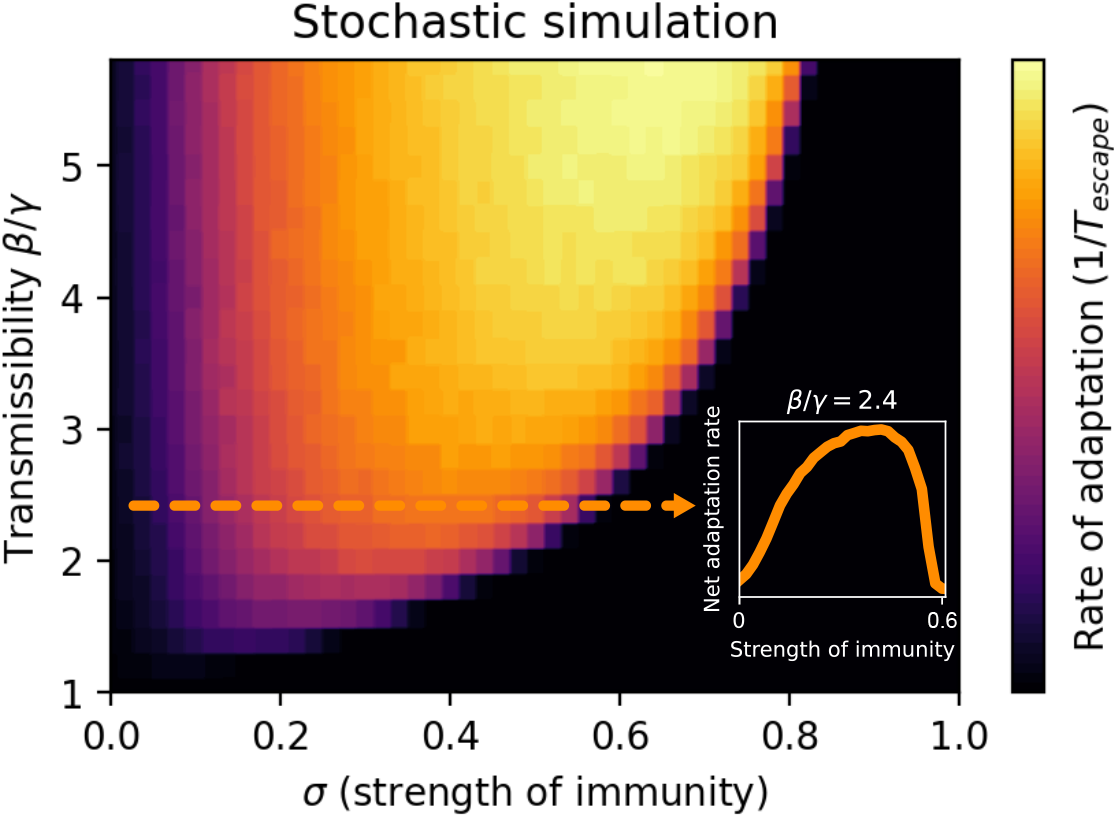
Stochastic (Doob-Gillespie) simulation of the Complete Exposure scenario. Note that this explicitly stochastic realization is qualitatively very similar to the analytic results presented in Fig. 2A of the main text.

## Notes

### Competing Interest Statement

The authors have declared no competing interest.

